# Assigning a social status from face adornments: an fMRI study

**DOI:** 10.1101/2023.03.08.531503

**Authors:** M Salagnon, F d’Errico, S Rigaud, E Mellet

## Abstract

The human face has been culturally modified for at least 150,000 years using practices like painting, tattooing and scarification to convey symbolic meanings and individual identity. The present study used functional magnetic resonance imaging to explore the brain networks involved in attributing social status from face decorations. Results showed the fusiform gyrus, orbitofrontal cortex, and salience network were involved in social encoding, categorization, and evaluation. The hippocampus and parahippocampus were activated due to the memory and associative skills required for the task, while the inferior frontal gyrus likely interpreted face ornaments as symbols. Resting-state functional connectivity analysis clarified the interaction between these regions. The study highlights the importance of these neural interactions in the symbolic interpretation of social markers on the human face, which were likely active in early Homo species and intensified with Homo sapiens populations as more complex technologies were developed to culturalize the human face.

## 1. Introduction

The use of technologies to change the appearance of our bodies to communicate information about our identity and social role dates back hundreds of thousands of years. Body painting, tattooing, scarification, wearing of ornaments, mutilations, hairstyles, and clothing are amongst the best-known practices for performing these functions in traditional societies ^1–5^. Personal ornaments, in particular, play a crucial role in communicating ethnic affiliation, reinforcing the sense of belonging to the group and its cohesion, establishing boundaries with neighboring groups, and conveying information on linguistic, ideological, and religious membership ^6–14^. Ornaments can also provide information about social status, gender, marital situation, and the number of children the wearer has had. Special ornaments and body paints may be put on at rites of passage occurring at the individual birth, during initiation ceremonies, marriage, healing, or death ^15–19^.

The earliest use of red ochre goes back to 500 ka in Africa ^20–23^, 380 ka in Europe ^22,24,25^, and 73 ka in Asia ^22,26,27^. Dapschauskas and colleagues (2022) identified three phases of ochre use in the African Middle Stone Age: an initial phase from 500 ka to 330 ka, an “emergent” phase from 330 ka to 160 ka, and a “habitual” phase from 160 ka to 40 ka. The latter phase, when a third of archaeological sites contain ochre, is interpreted by these authors as the manifestation of intensifying ritual activity in early populations of *Homo sapiens*. This view is consistent with the results of studies indicating that in this last and the previous phase, certain types of mineral pigments were transported over long distances ^20,28,29^, that certain shades of red were particularly sought after ^30–32^, that ochre was modified by heating to change its color ^33,34^ but see ^35^, and that in some cases very small quantities of pigments were produced ^36^, a behavior more consistent with a symbolic than a utilitarian function.

The wearing of personal ornaments, many of which are deliberately covered with ochre, is attested since at least 142 ka in North Africa, 80 ka in Southern Africa, and 120 ka in the Near East ^37–45^. Because the understanding by others of the meaning attached to ornaments and body paints presupposes the existence of shared codes, archaeological objects which have fulfilled these functions are often considered reliable evidence for the emergence of language and symbolic material cultures in our genus ^40,41,46–49^. In this regard, wearing body adornments can be considered an archaeological indicator of modern social cognition.

Although body symbols played a key role in all human societies and appeared very early in human history, the cerebral regions mobilized by their perception and interpretation remain unknown. Numerous studies have focused on the brain substrates of the emotional aspects of social cognition and perspective-taking (Theory of Mind). They have emphasized the role of the medial prefrontal cortex, the temporoparietal junction, and the temporal poles ^50,51^. However, one study showed that social status recognition was minimally disrupted following ventromedial lesions, suggesting that the network involved in this function would be distinct from that dealing with the emotional aspects of social cognition ^52^. Neuroimaging studies have confirmed that the perception of social hierarchies relies on the intraparietal sulcus, the dorsolateral and orbital frontal cortex, and the lateral occipital and occipitotemporal cortex ^53–56^. However, the identification of social markers does not necessarily imply a ranking.

Body ornaments and facial paintings may convey information on social roles disconnected from a social hierarchy. Although body paintings and the wearing of beads to express social roles are attested in the earliest *Homo sapiens* and probably in Neanderthals ^57–59^, very little is known about the brain networks involved in processing such information, the possible processes that led to a complexification of these behaviors, as well as their timeline. In the present study, participants were asked to assign social roles or statuses to faces adorned with paintings, beads, or both. At the same time, their brain activity was monitored using functional magnetic resonance imaging (fMRI). Participants were given no guidance on the meaning of the face decorations and had to create their arbitrary social code. The attribution of a social status mobilizes implicit and explicit processes. Implicit processes are rapid, require little cognitive effort, and can occur without awareness. Explicit processes are cognitively demanding, slow, and deliberative ^60^. To isolate explicit processes, we included, using the same stimuli, a perceptual task (1-back) that does not explicitly require a social role attribution. Brain activity during this task was compared to that performed during explicit social status attributions. In addition, at rest, the functional connectivity of the brain regions involved was analyzed to provide information on the interaction of the brain areas implicated in the social status attribution task. Our results identify, for the first time, the brain networks engaged in attributing social status from different arrangements of paintings and ornaments on the human face, the way they work in synergy, and provide sound bases on which build an evolutionary scenario for the gradual integration of these brain areas during the evolution of our genus.

## 2. Materials and methods

### 2.1. Ethics statements

The ‘Nord-Ouest III’ local Ethics Committee approved the study on 10/14/2021 (N°IDRCB: 2021-A01817-34). All the participants signed informed consent before the MRI acquisition.

### 2.2. Participants

Thirty-five healthy adults (age range 18–29 years, mean age 22 ± 2 years (SD), 18 women, four left-handed) with no neurological history were included. One participant was excluded from the analysis because of a brain abnormality discovered during MRI acquisition.

### 2.3. Experimental design

The functional acquisition was organized in a single session consisting of six runs during which participants had to perform a selection task (first three runs), then a 1-back task (last three runs). After receiving instructions for these tasks, participants completed a short training run outside the MRI.

#### 2.3.1. Stimuli

The set of stimuli included pictures of faces (up to below the shoulders) of 34 unknown people in the same range of age (17 women, 17 men, around 30 years old) wearing ornaments and adopting a neutral expression. Each face was ornamented with either spherical wooden beads, red paintings, or a combination of both (Figure 1). Ornaments included earrings with one or three beads, necklaces with one or two chains of beads, a diadem consisting of a chain of beads, and a single large spherical bead in the middle of the forehead. Red paintings included one or three vertical lines on the chin; a dot or a horizontal line on the forehead; oblique lines on the cheeks; and a large horizontal band including the eyes. Associations of paintings and beads were designed to make both types of ornamentation gradually more invasive on the face. In all, twelve types of facial ornamentation have been designed and implemented.

**Figure 1.**
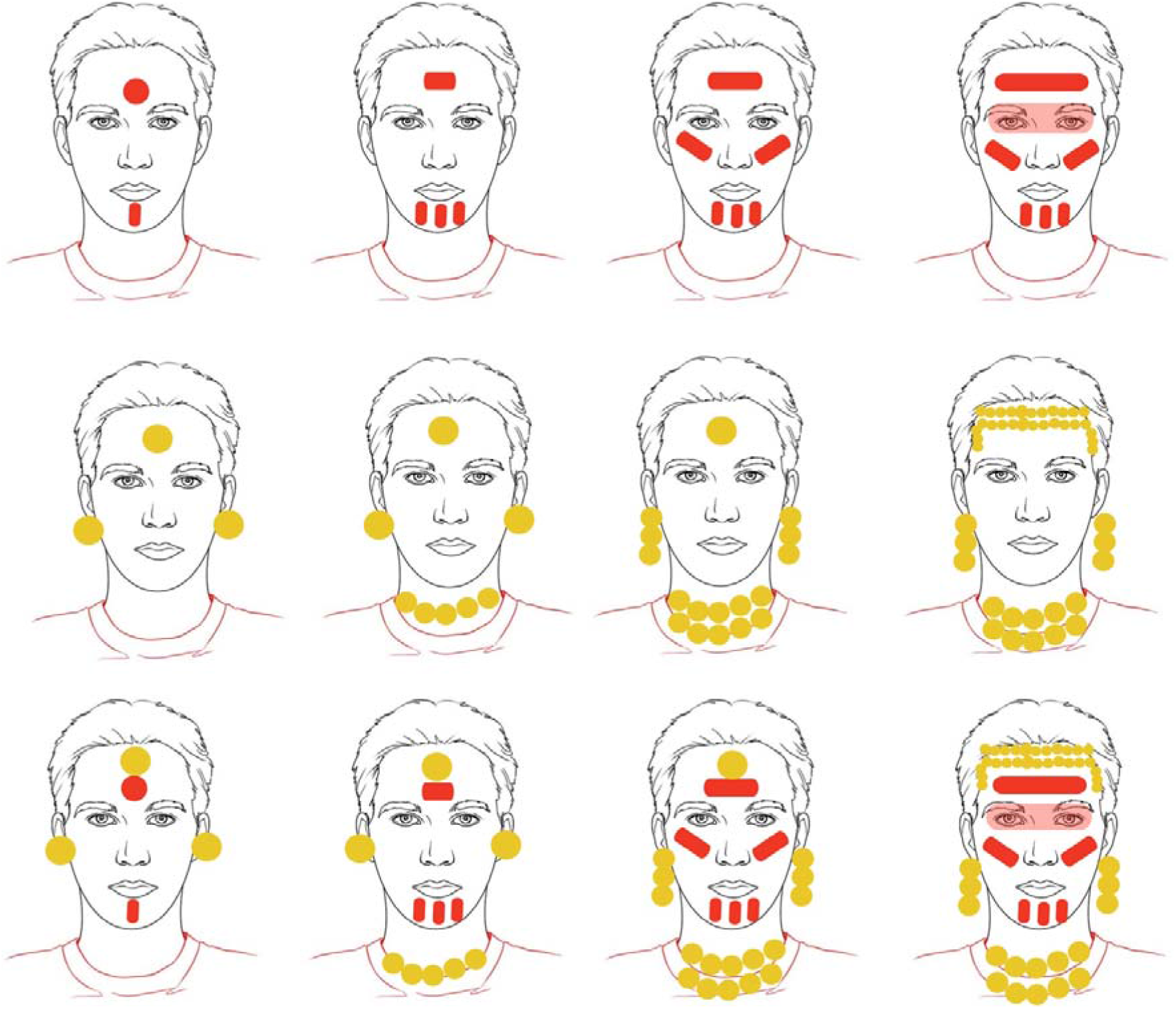
Face ornamentations used in the tasks. Top row: paint only. The middle row: beadworks only. Bottom row: a combination of paint and beadworks.

#### 2.3.2. Selection task (event-related paradigm)

These three runs followed a slow event-related design, i.e., the change in the BOLD signal was collected for each stimulus presentation, and the time between each presentation allowed the signal to return to its baseline level. The order of presentation was randomized. The stimuli corresponded to a triplet of photos of three different persons of the same gender (male or female), one wearing ornaments, one with paintings, and one with both (Figure 2). Within a triplet, the richness of the ornaments was comparable between the pictures to avoid biases in the choice. There were three levels of richness between the triplets (Figure 2).

**Figure 2.**
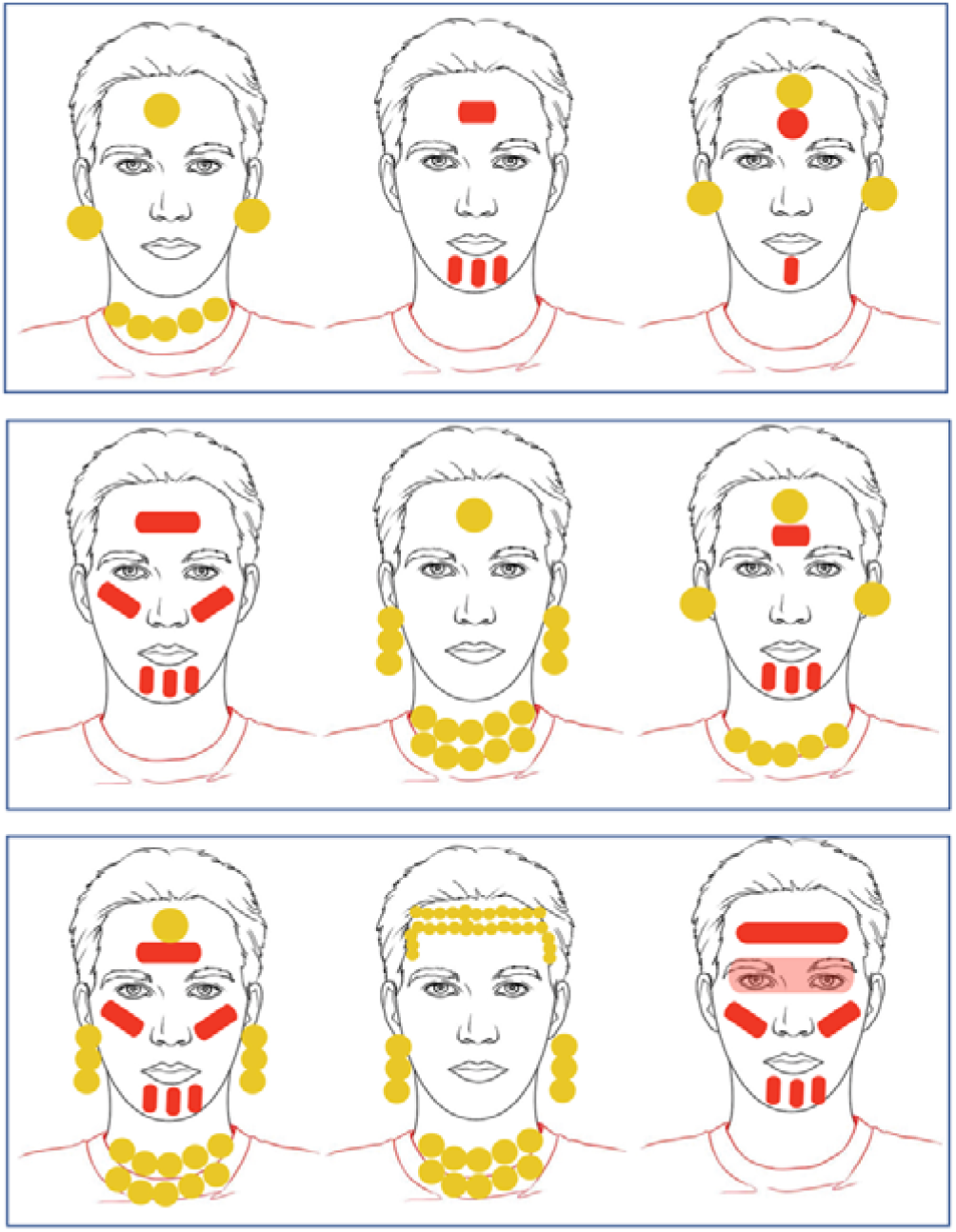
Drawn version of some triplets of stimuli used in the selection task. Each triplet was composed of individuals of the same sex (male or female). The three ranks correspond to three levels of richness. Note that in the study, photographs of real people with adorned face were presented but could not be displayed here because of their identifying nature.

The selection task was implemented as follows (Figure 3): a question was displayed during 0.75 s. The question could concern either the displayed persons’ social role (Social status condition) or the type of ornamentation they displayed (Ornament check condition). Then, a new triplet of pictures was shown for 4 s. The participant had to choose, by pressing the corresponding button of a response box as soon as they made their decision, the person who best fitted the proposed social status, e.g., “Shaman” (Social status condition) or the person who corresponded to the ornament type proposition, e.g., “Painted cheeks” (Ornament check condition). Each question was asked twice for each gender. The list of questions is displayed in Table 1.

**Figure 3.**
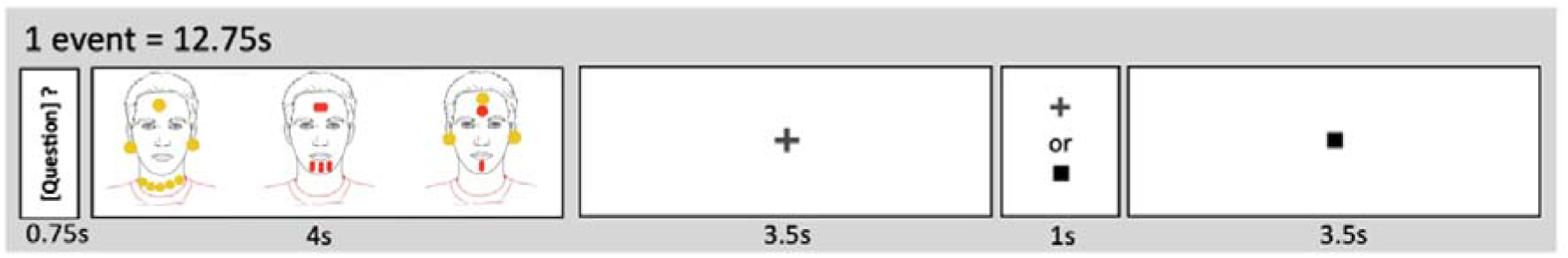
Organization of one event of the selection task.

**Table 1.**
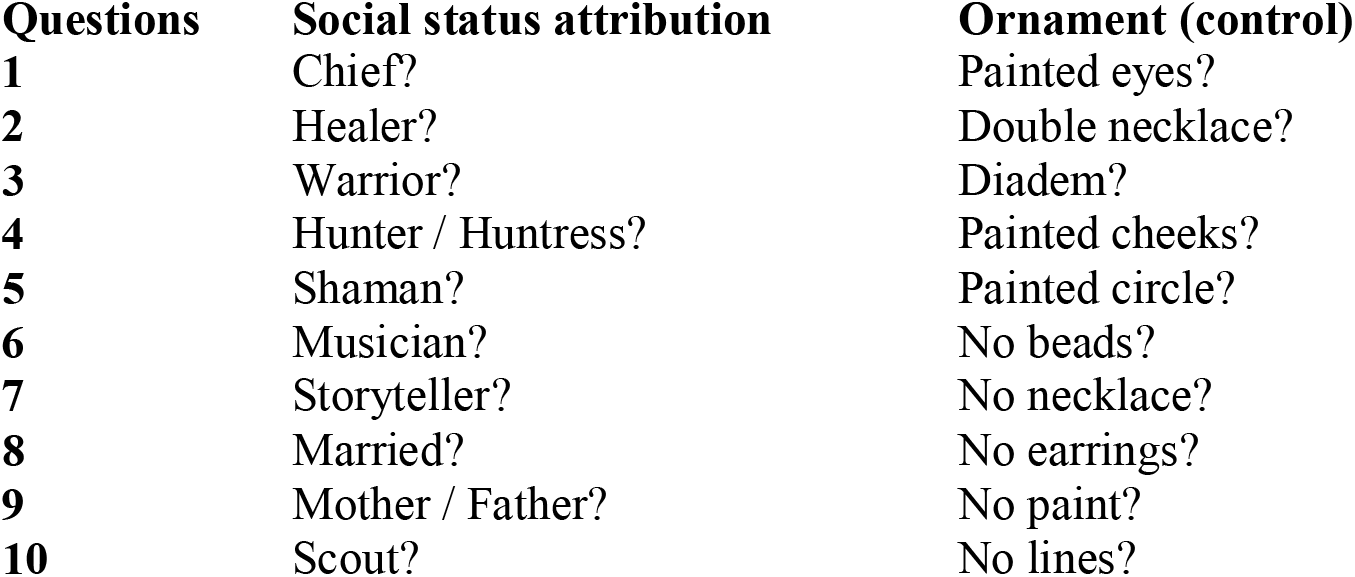
Questions in the social status and ornament check conditions. The wording of the questions was gendered according to the stimulus.

| Questions | Social status attribution | Ornament (control) |
| --- | --- | --- |
| 1 | Chief? | Painted eyes? |
| 2 | Healer? | Double necklace? |
| 3 | Warrior? | Diadem? |
| 4 | Hunter / Huntress? | Painted cheeks? |
| 5 | Shaman? | Painted circle? |
| 6 | Musician? | No beads? |
| 7 | Storyteller? | No necklace? |
| 8 | Married? | No earrings? |
| 9 | Mother / Father? | No paint? |
| 10 | Scout? | No lines? |

The Ornament check condition was designed as a control condition, bearing the same pictures as the social status condition. It required attention to the ornaments without implementing social cognition processes. Then, a fixation cross was displayed, and a square appeared after a variable delay (3.5 s ± 1 s). Participants had to click the “1” button on the response box when the square appeared. This constituted the baseline, allowing the BOLD signal to return to its baseline level between events. Each event lasted 12.75 s.

Over the three runs, participants saw 80 stimuli, 40 in the social status condition and 40 in the ornament check condition. Each run lasted 5 min and 51 s each and included 27 events (except run C, which included one less event) for a total duration of 5 min 38 s). Stimuli were presented in random order within each run. Immediately after the MRI acquisition, the experimenter asked the participant the criteria on which they based their social role attribution in the status condition.

#### 2.3.3. 1-back task (block design)

In the 1-back task, participants viewed a succession of ornamented faces (displayed for 1 s each, with an interstimulus interval of 983 ms). The participants had to report the repetition of two faces (Face condition) or two types of ornamentation (Ornament condition, including three modalities: paintings, beads, or both simultaneously) by pressing the “1” button on the response box. This repetition criterion was displayed during 750 ms at the beginning of each block. Fifteen stimuli belonging to the same category of ornamentation were presented within the same block (i.e., within a block, there were no images belonging to different categories). There were three repetitions per block. Each of the three runs lasted 4 min and 15 s and included six experimental blocks of 30.6 s interspersed with seven fixation blocks of 10.2 s. Each run had four blocks of ornament condition and two blocks of face condition. The presentation order of the 1-back runs was randomized.

### 2.4. MRI acquisition

Neuroimaging data acquisition was performed using a Siemens Prisma 3 Tesla MRI scanner. Structural images were acquired using a high-resolution T1-weighted 3D sequence (TR = 2000 ms, TE = 2.03 ms; flip angle = 8°; 192 slices and isotropic voxel volume of 1 mm^3^). Functional images were obtained using a whole-brain T2*-weighted echo planar image acquisition (T2*-EPI Multiband x6, sequence parameters: TR = 850 ms; TE = 35 ms; flip angle = 56°; 66 axial slices and isotropic voxel size of 2.4 mm^3^). The first sequence lasted 8 min and recorded participants’ brain activity during resting state (i.e., when they let their thoughts flow freely, without having a task to perform or falling asleep). This acquisition was used to perform a resting-state functional connectivity analysis. Then, functional images were acquired when the participants performed tasks based on stimuli perception. This was done during six runs (three for each task: selection and 1-back). The presentation of the experiment was programmed in E-prime software 3.0 (Psychology Software Tools, Pittsburgh, PA, USA). The stimuli were displayed on a 27” screen. Participants saw the stimuli through the back of the magnet tunnel via a mirror mounted on the head antenna.

### 2.5. Data analysis

#### 2.5.1. Behavioral analysis

For the selection task, we evaluated the effects of condition (Social status or Ornament check), participant gender, and stimulus gender on reaction time using a linear mixed-effects model, adjusting for random effects at the participant level. A three-factor interaction term between condition, participant gender, and stimulus gender (and their lower-order terms) was defined as fixed-effect predictors and reaction time as the dependent variable. The significance of fixed effects was assessed through ANOVA components.

#### 2.5.2. Functional neuroimaging analysis

T1-weighted scans were normalized via a specific template (T1-80TVS) corresponding to the MNI space using SPM12. The 192 EPI-BOLD scans were realigned in each run using a rigid transformation to correct the participant’s motion during the fMRI sessions. Then, the EPI-BOLD scans were rigidly registered structurally to the T1-weighted scan. All registration matrices were combined to warp the EPI-BOLD functional scans to standard space with trilinear interpolation. Once in standard space, a 5-mm-wavelength Gaussian filter was applied.

In the first level analysis, a generalized linear model (GLM, statistical parametric mapping (SPM 12), http://www.fil.ion.ucl.ac.uk/spm/) was performed for each participant to process the task-related fMRI data, with the effects of interest (tasks) modeled by boxcar functions corresponding to events or blocks, convolved with the standard hemodynamic SPM temporal response function. We then calculated the effect of individual contrast maps corresponding to each experimental condition. Note that eight non-interest regressors were included in the GLM analysis: time series for white matter, CSF (average time series of voxels belonging to each tissue class), the six motion parameters, and linear temporal drift.

Group analysis (second-level analysis) of fMRI data was conducted using JMP^®^ software, version 15. SAS Institute Inc, Cary, NC, 1989-2019. The first step was to select the brain regions activated in the contrasts of interest, namely [Social status minus Ornament check] in the selection task and [Ornament minus Face] in the 1-back task. We extracted signal values from the [Social status minus Ornament check] contrast from each brain region of each participant (hROI, homotopic region of interest) in the AICHA atlas ^61^. The MNI coordinates of the center of mass of each activated hROIs are given in the supplementary material section. The hROIs included in the analysis fulfilled the following criteria: significantly activated in the [Social status minus Ornament check] contrast (univariate t-test p < 0.05 FDR corrected); and significantly activated in the [Social status minus baseline] contrast (univariate t-test, p < 0.1 uncorrected) to eliminate deactivated hROIs. 32 regions whose BOLD signal occupancy was less than 80% (susceptibility artifacts) were excluded from the analysis. The hROIs excluded are listed in the supplementary material.

This procedure led to 95 hROIs being more activated in the social status condition than in the ornament check condition. The same method was applied to the [Ornament minus Face] contrast and [Ornament minus baseline] contrast leading to 81 activated hROIS for the 1-back task. In addition, we applied a univariate t-test (FDR corrected, p < 0.05) to compare the BOLD values in the 95 hROIs activated in the [Social status minus Ornament check] contrast to those 81 hROIs elicited by the [Ornament minus Face] contrast of the 1-back task. This allowed for refining the specificity of the regions involved in the social status attribution and its explicit components. Thirty-seven hROIs were more activated in the [Social status minus Ornament check] contrast than in the [Ornament minus Face] contrast.

#### 2.5.3. Resting-state analysis

The task-based functional analysis was complemented with a resting-state functional connectivity analysis using the CONN v 20.b toolbox software ^62^, which runs under MATLAB 2021a.

Functional imaging data were pre-processed using the CONN default pre-processing pipeline for volume-based analyses. The steps for functional data comprise realignment and unwarping for subject motion estimation and correction (12 parameters). Next, centering to (0,0,0) coordinates and ART-based outlier detection identification was applied. Segmentation and normalization to MNI space were applied next. Structural data were translated to (0,0,0) center coordinates, segmented (gray/white/CSF), and normalized to MNI space. In the denoising step, we applied band-pass filtering (0.01–0.1 Hz) after regression of realignment parameters (12), white and gray matter, and CSF confounds. Then, we applied linear detrending and despiking after regression. For the ROI to ROI functional connectivity analyses, we used AICHA atlas ^61^. We considered the 95 hROIs activated in the [Status minus Ornament] contrast. For group-level results, we calculated ROI-to-ROI connectivity correlations, threshold with a unilateral t-test, and FDR-corrected p < 0.05.

## 3. Results

### 3.1. Behavioral results

Participants responded faster in the ornament check condition (mean response time ± SD: 1.3s ± 0.5s) than in the social status condition (mean response time ± SD: 2s ± 0.8s): *F*_(1,32)_ = 227.8 p < 0.0001. Participant gender and stimulus gender had no significant effects (either main or interactions).

### 3.2. Post-MRI debriefing of the selection task

Twenty-two participants reported that they considered ornamentation a more important criterion than phenotype in assigning a social role/status. A few reported they sometimes paid attention to facial features, for example, in cases of indecision or for specific roles such as father/mother. Eleven participants reported paying more attention to facial characteristics than to ornamentation. Two participants stated that the most important criterion for them (facial features or ornamentation) varied according to the questions. All participants reported that they never answered randomly, except in rare exceptions. Participants generally reported having an attribution strategy in place that they maintained throughout the experiment. For example, some participants associated the absence of beads with a mobile role, such as scout or hunter. For the same role, there was not necessarily a consensus among participants. For example, some participants attributed warrior status to faces wearing only beads, while others attributed this status to faces bearing only paintings.

### 3.3. Neuroimaging results

#### 3.3.1. Social status minus Ornament check (event-related paradigm)

The [Social status minus Ornament check] contrast revealed a set of 95 cortical and subcortical regions that were more activated when participants assigned social status to adorned faces than when they assessed the type of ornamentation (Table 2, Figure 4).

**Table 2.**
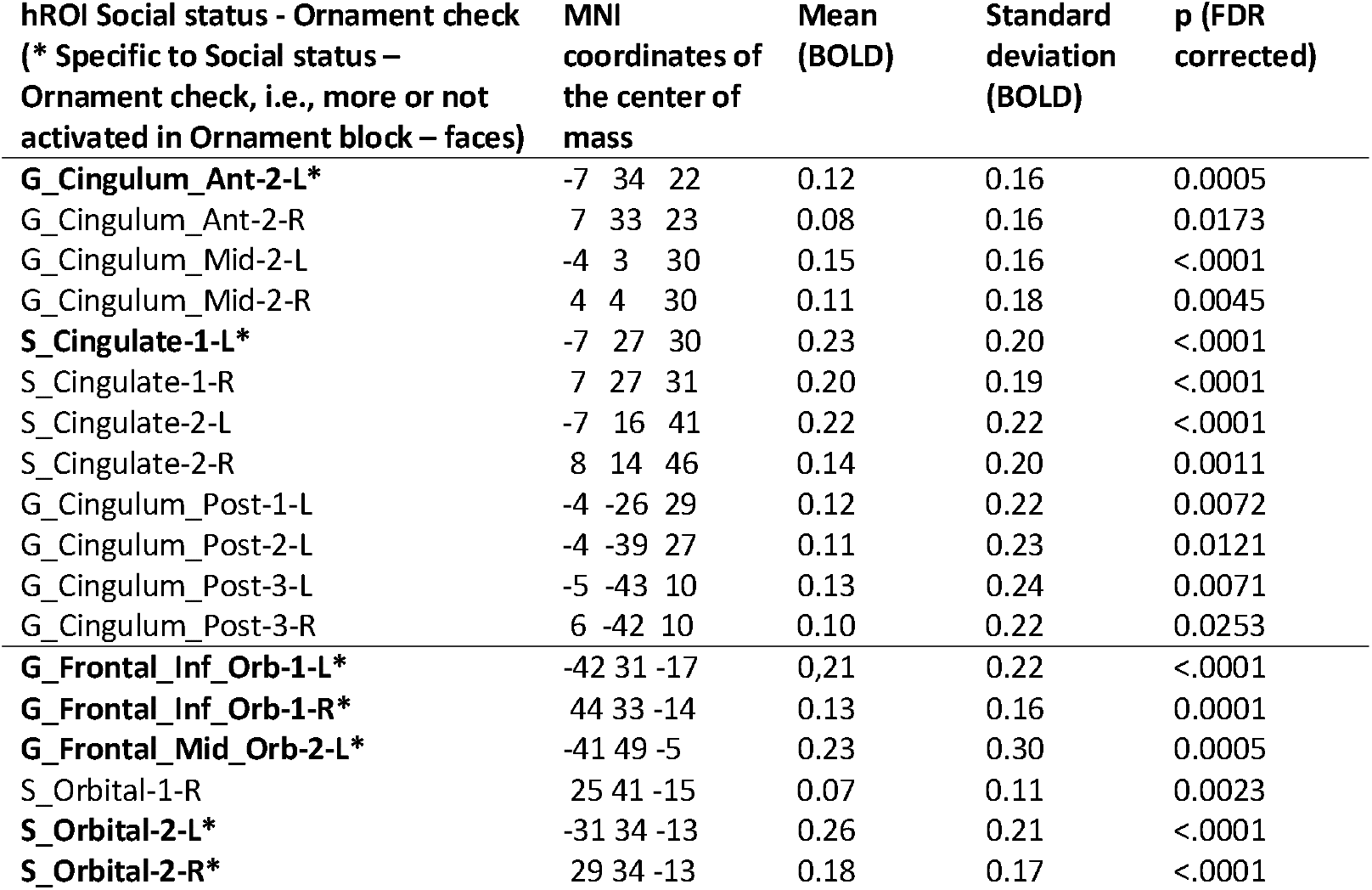
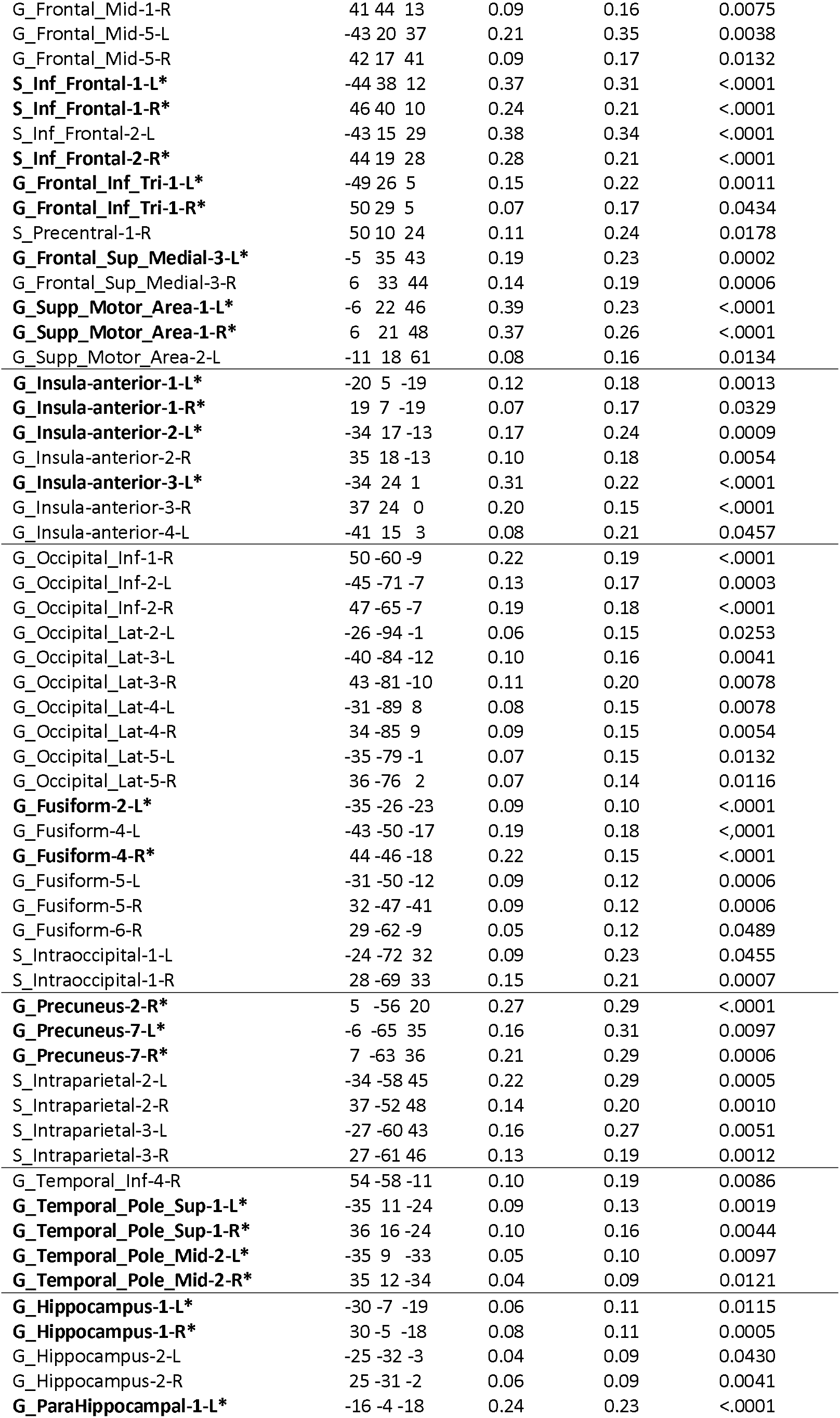
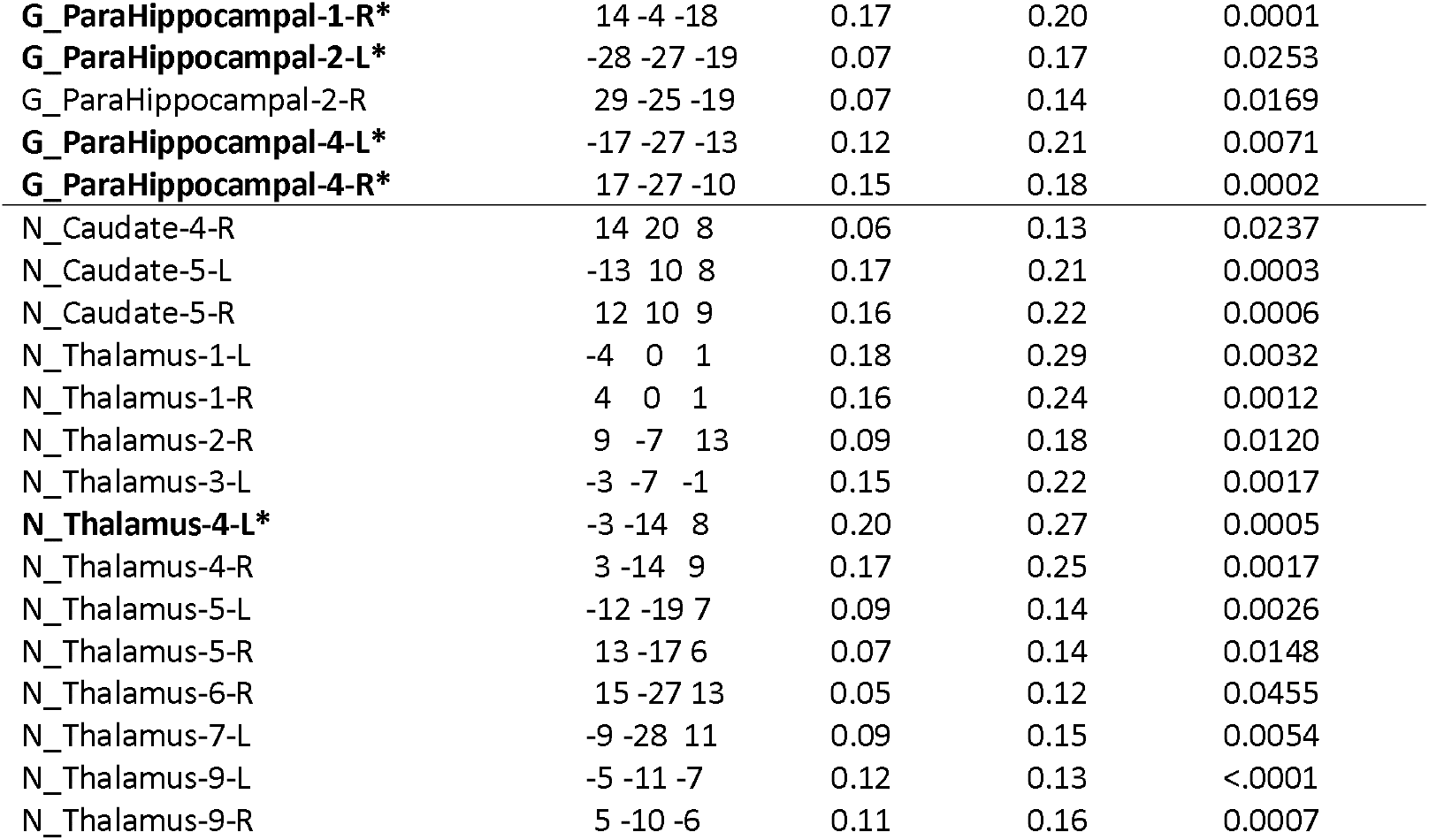
Mean, standard deviation, and p-value of the activated regions in the [Social status minus Ornament check] contrast.

**Figure 4.**
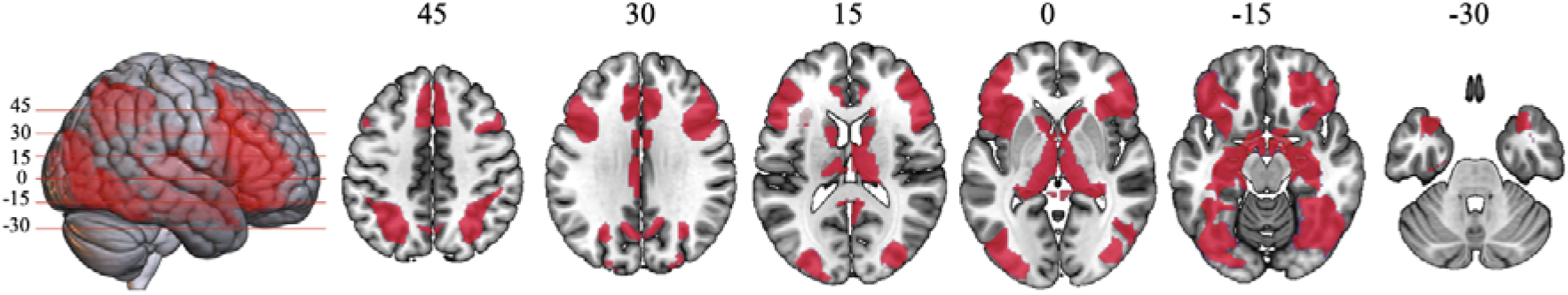
Activated regions in [Social status minus Ornament check] contrast superimposed on an MRI template. Numbers indicate the z value of the axial slice in MNI space.

In the occipital lobe, these regions included the lateral occipital cortex and the fusiform gyrus (including the Fusiform Face Area, FFA). We used the Neurosynth platform ^63^ to synthesize the activations reported in the literature during face perception and ensure their consistency with our results. We conducted a meta-analysis including 125 studies that contained the term “neutral face” in their abstracts, i.e., pictures of faces adopting a neutral expression. It evidenced the involvement of a right fusiform region (MNI coordinates of the activation peak: 38, −42, −16). This matched the location of the G_Fusiform-4-R in our AICHA atlas (MNI coordinates of the center of mass: 44, −46, −18). The FFA occupied a large portion of this functional region of the AICHA atlas.

The activations extended to the parahippocampal gyrus on the medial side of the temporal lobe. In the parietal lobe, the intraparietal sulcus was activated bilaterally. In the frontal lobe, activations included the middle and inferior frontal gyri on the lateral side and the anterior part of the supplementary motor area medially. Activations also concerned several paralimbic and limbic cortex regions, such as the anterior insula, the anterior cingulate, the posterior cingulate and adjacent precuneus, the orbitofrontal cortex, the temporal poles, and the hippocampus. The subcortical structures, the head of the caudate nucleus, and the thalamus, especially in its mediodorsal part, were also involved.

#### 3.3.2. Ornament minus Face (1-back paradigm)

The [Ornament minus Face] contrast revealed a set of 81 cortical and subcortical regions, which were more activated when participants checked the repetition of ornamentation than when they looked for the repetition of faces. These regions were mostly located in the lateral and inferior occipital cortices and the fusiform gyrus, extending to the inferior temporal gyrus. Participants also activated the intraparietal sulcus, the anterior insula, and some frontal regions, such as the superior frontal sulcus, inferior frontal sulcus, the supplementary area, and the middle frontal gyrus.

Among these regions, 37 hROIs were significantly less activated in the [Ornament minus Face] contrast than in the [Social status minus Ornament check] contrast (Table 2).

#### 3.3.3. Resting-state functional connectivity

The resting-state functional connectivity analysis revealed 348 positive connections significant across subjects (p < 0.05 FDR, univariate t-test) between the 37 hROIs. T-values varied from 2 to 25 (Figure 5). These 37 hROIs can be divided into two groups based on their resting-state functional connectivity. A network connected the precuneus and temporal lobe regions, including the hippocampus, the parahippocampal cortex, the temporal pole, and a part of the fusiform gyrus. A second network connected mainly frontal regions, including the inferior frontal sulcus and gyrus, the orbitofrontal cortex, the dorsal anterior cingulate cortex, the supplementary motor area, and the anterior insula.

**Figure 5.**
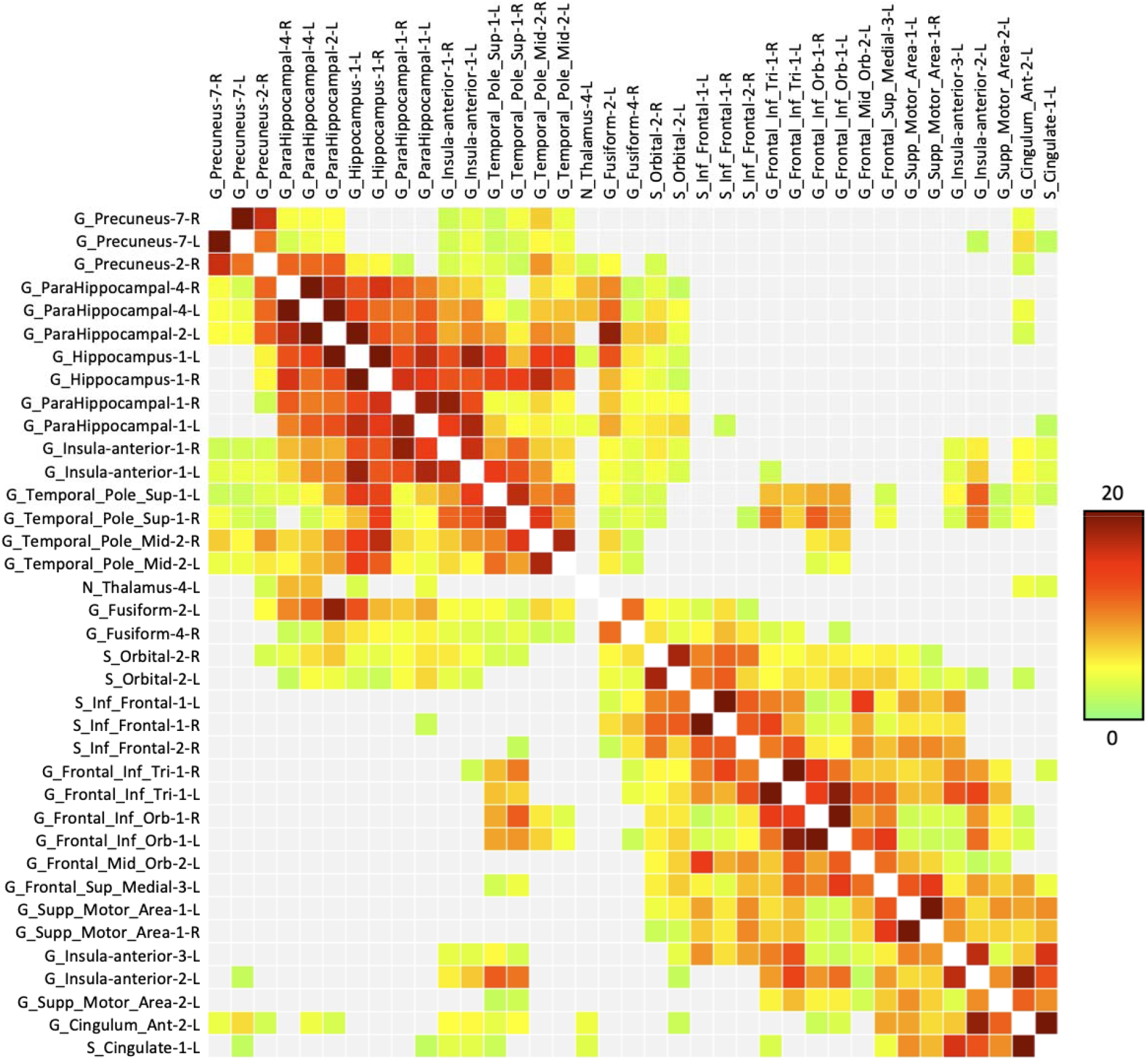
Resting-state connectivity matrix of 37 hROIs specific to the social status assignment (i.e., positive in the contrast [Social status minus Ornament check] and positive in contrast [Ornament (1-back) minus Faces]. The color scale (green to red) reflects the t-value on each connection averaged across subjects.

The G-Fusiform-4-R and the Temporal_Pole_Sup-1-R regions were connected to 22 and 27 hROIS, respectively. The G-Fusiform-4-R and the Temporal_Pole_Sup-1-R regions were strongly connected to their group and many regions of the other group (see supplementary materials for detailed results). The S_Orbital-2 was connected to 26 hROIs.

## 4. Discussion

This study aimed to identify the brain regions involved in attributing social status from the visual analysis of adorned faces. Adorning one’s body to transmit social information represents a symbolic behavior that appeared at least 150,000 years ago and probably much earlier. Therefore, we can assume that the networks revealed in the present study were, at least to a degree, functional in the earliest *Homo sapiens* and contemporary or earlier hominins displaying such behaviors.

These regions can be categorized into four groups: 1. occipitotemporal regions of the ventral visual pathway, including lateral occipital regions, fusiform gyrus, parahippocampal gyrus extending to the hippocampus, and the temporal poles; 2. regions belonging to the salience network such as the anterior insula and the anterior cingulate cortex; 3. the intraparietal sulcus; and 4. the ventral and dorsal regions of the lateral prefrontal cortex and the orbitofrontal cortex.

Some of these regions were also activated in the 1-back task, indicating that they are not specific to an explicit social attribution but may be involved in an implicit social appraisal. This is the case for most visual regions (except Fusiform-4-R and Fusiform-2-L), the intraparietal sulcus, and most of the thalamus and retrosplenial regions. In contrast, activity in the inferior and orbital frontal areas, hippocampal and parahippocampal regions, the temporal poles, and the salience network, including the anterior cingulate and parts of the anterior insula, remained significant when activity in these regions during the 1-back task was subtracted.

### 4.1. Visual Ventral pathway and medial temporal regions

Lateral and ventral occipital regions were more activated by the social status attribution than by the assessment of decoration type. This suggests that deeper visual processing is required to attribute a social status. Most of these occipitotemporal regions were also activated during the 1-back task and were thus not specifically involved in assigning a social status to adorned faces. However, two hROIs were significantly more activated during social status attribution than in the 1-back task, namely G_Fusiform-2-L and G_Fusiform-4-R. The latter is particularly interesting since it includes the so-called fusiform face area (FFA), which is sensitive to face perception ^64–66^ and lateralized in the right hemisphere ^67^. Thus, although all conditions included face perception and none required specific attention to faces, FFA appeared more solicited by social status assignment. It has been shown that FFA is sensitive to physical characteristics and their possible social correlates ^68,69^. More recently, a study showed that FFA processes characteristics such as social traits, gender, and high-level visual features of faces ^70^ and might thus initiate the social processing of faces. The results of the present study suggest that, in the context of social role attribution, FFA can process non-physiognomic features. This is consistent with the fact that the FFA promotes holistic rather than local processing ^71–73^. Ornamented faces may have been perceived as a whole in the social attribution task, while attention was focused on details during the assessment of decoration type and the 1-back task. In other words, attributing social status involves a more complex process relying on a set of components, such as the types of decoration, their association, their location on the face, and the face itself.

In summary, the activation of FFA in our social status assignment task could reflect the implementation of preliminary social categorization processes based on a holistic analysis of ornamented faces, which is further achieved in other regions of the brain, particularly the orbitofrontal cortex.

In the anterior extension of the ventral visual pathway, we found that the hippocampus and parahippocampal gyrus were more activated by the social status attribution than by the ornament type assignment and significantly more activated when compared to the 1-back task, reflecting their specificity to social status attribution. The hippocampus reflects episodic memorization processes strongly involved in social cognition ^74,75^. The parahippocampus appears to play, among others, a pivotal role in contextual associative processing ^76,77^, i.e., in binding elements composing stimuli. It provides a unified context for further processing (see ^78^ for a review). In the framework of the present study, participants arbitrarily associated face decorations with social status. After the fMRI sessions, they reported that once they had established an ornament/status association strategy, they stuck to it throughout the sessions, with exceptional random responses. Contextual associations were thus an essential aspect of the processes involved in the status assignment task. The activation of the parahippocampal cortex reasonably reflects the implementation of these processes. It has been suggested that the anterior part of the parahippocampus preferentially processes non-spatial contextual associations, and the posterior part, comprising the parahippocampal Place Area (PPA), spatial associations ^76,79^. In the present study, the activation of the anterior parahippocampus is consistent with the non-spatial nature of the associations.

The activation of the medial temporal gyrus might be linked to one of the temporal poles. Several studies have documented the involvement of the temporal pole in social cognition, and this region is considered part of the social brain network ^80–84^. Although its role is still under discussion, it has been proposed that this brain area is involved in encoding and retrieving social knowledge ^85^. As was the case in this study, assigning social status mobilizes stereotypical social knowledge (e.g., the chief must have the most ornaments) and entails encoding: The participants associated a type of ornamentation with a social role and created an arbitrary social code that they reused throughout the task. Thus, we propose that the parahippocampus and the temporal pole, which are strongly functionally connected, work in synergy to facilitate the association of a type of ornamentation with a specific social status and then to encode and restore this association.

### 4.2. Inferior and orbitofrontal cortex

Assigning a social status involved many frontal regions not solicited during the ornament type attribution condition. However, the specific areas for explicit processes, i.e., activated in the social attribution task compared to the 1-back task, were mainly in the lateral part of the inferior frontal gyrus and the orbitofrontal cortex as defined by Rudebeck and Rich ^86^. The resting-state connectivity analysis showed that these regions were highly functionally linked. Previous studies have emphasized the role of the orbitofrontal cortex in social cognition in non-human primates and humans. It has been argued that this cortical area contains neurons sensitive to representing social categories ^87^ and evaluating social information ^88^ in non-human primates. In humans, a deficit in social perception after orbitofrontal cortex lesions ^89^, an inability to judge social traits in a decision-making task ^90^, or acquired sociopathy have been reported ^91^.

In healthy participants, fMRI studies have emphasized the role of the orbitofrontal cortex in social cognition and social behavior (^55,92^; See ^60^ for a review) and, more specifically, in explicit processing ^93^. The orbitofrontal cortex is sensitive to non-verbal social signals ^55^. Recent results indicate that this area is critical in representing social status ^92^. A recent fMRI study showed that the OFC represented the stereotypic social traits of others and that its pattern of activity was predictive of individual choices, highlighting its critical role in social decision-making ^94^. In these studies, participants had to behave according to the facial expression, attitude, or social category of the individuals presented in the experiment. Our results extend these findings. Unlike previous studies, participants based their decision on symbolic features (the type and arrangement of ornamentations), to which they arbitrarily attributed social meaning. This implies that the role of the orbitofrontal cortex in social decision-making is not restricted to processing stereotypical attitudes or social groups, a capacity shared with non-human primates. Social evaluation based on symbolic external attributes also involves this region in humans.

The social status attribution task heavily relies on high-order executive functions such as attentional control, selection, and flexibility. The activation of the pars triangularis of the inferior frontal gyrus extending to the inferior frontal sulcus reflects these aspects ^95^. Although the activation was bilateral, the right and left inferior frontal gyrus probably played a different role in the task. The right inferior frontal gyrus is explicitly associated with high-level social cognition ^96^. The left inferior frontal gyrus is involved in selecting some aspects or subsets of available information among competing alternatives ^97,98^. This region also plays a role in processing non-linguistic symbolic information ^99,100^, consistent with the symbolic value attributed by the participants to face adornments.

Overall, the prefrontal cortex’s involvement in the present study underlines its role in social decision-making. Our results extend their contribution to symbolic social communication, here materialized by face ornamentations.

### 4.3. Salience network

Social status attribution elicited activation in the anterior insula, the dorsal anterior cingulate cortex (dACC)/pre-SMA, and subcortical structures, such as the thalamus and the caudate nuclei. These regions constitute the so-called salience network, whose key components are the insula anterior and the dACC/pre-SMA ^101–103^. This network is involved in selecting relevant elements of the environment for perceptual decision-making ^104–106^. In our case, participants had to extract salient information from the ornamented faces to associate the proposed social status with one of the three faces presented to them. The salience network was also activated in the 1-back task by the need to detect the repetition of ornamental patterns. However, the greater uncertainty in decision-making during the attribution task can explain why activation of the salience network was more extensive during status attribution than the 1-back task ^107^. To attribute a social status, participants had to make a forced choice among three possibilities, with several plausible answers and had to compare the different options and arbitrate to choose only one. These aspects of the task have probably triggered the activation of the dACC/pre-SMA, belonging to the salience network. The dACC/pre-SMA has been reported as involved in conflict and performance monitoring ^108–110^, and more recently in social categorization domain ^111^.

### 4.4. Resting-state functional connectivity

Resting-state functional connectivity provides insight into the potential interactions between neural assemblies activated by the social status assignment task. The G_Fusiform-4-R and the G_Temporal_Pole_Sup-1-R were characterized by many connections with other activated regions (Figure 6). These two regions were connected with 22 and 27 hROIs, respectively. The G_Fusiform-4-R included the FFA (see results) and is likely involved in the initial processing phase. The functional relationships between the medial temporal lobe, the fusiform gyrus, and the temporal pole reflected the association of the perceptive, social, mnemonic, and associative aspects of the task. In addition, the G_Temporal_Pole_Sup-1-R was connected with frontal regions and could act as a hub, allowing communication between visual areas and executive frontal regions. The connection between the temporal pole and the salience network enables the exchange of information necessary for evaluating and selecting inputs relevant to social decision-making. The temporal pole and the salience network were related to the inferior frontal gyrus, contributing to the evaluation of subjective confidence about a perceptual decision ^112^. The orbitofrontal cortex was functionally connected to the G_Temporal_Pole_Sup-1-R and the G_Fusiform-4-R. These regions, whose essential role in social status evaluation has been discussed above, could constitute the core network in the social attribution task (Figure 6). They must have allowed the integration of information leading to the assignment of social status based on the perception of symbolic cues.

**Figure 6.**
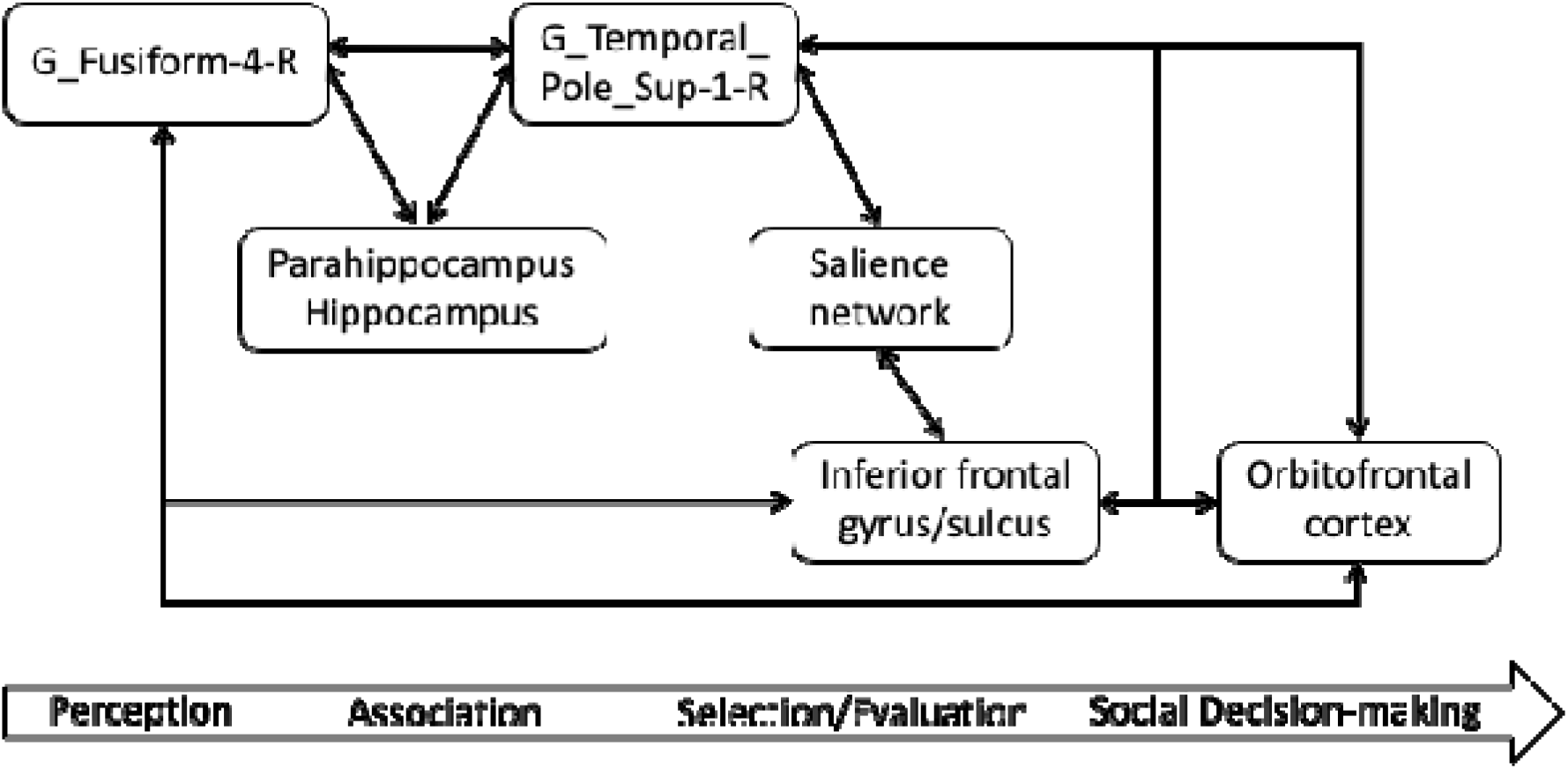
Schematic resting-state functional connectivity network between regions activated during a social status attribution task based on symbolic culturalized faces. Black arrows indicate the reciprocal resting-state functional connectivity between brain regions (univariate t-test, p < 0.05, FDR corrected).

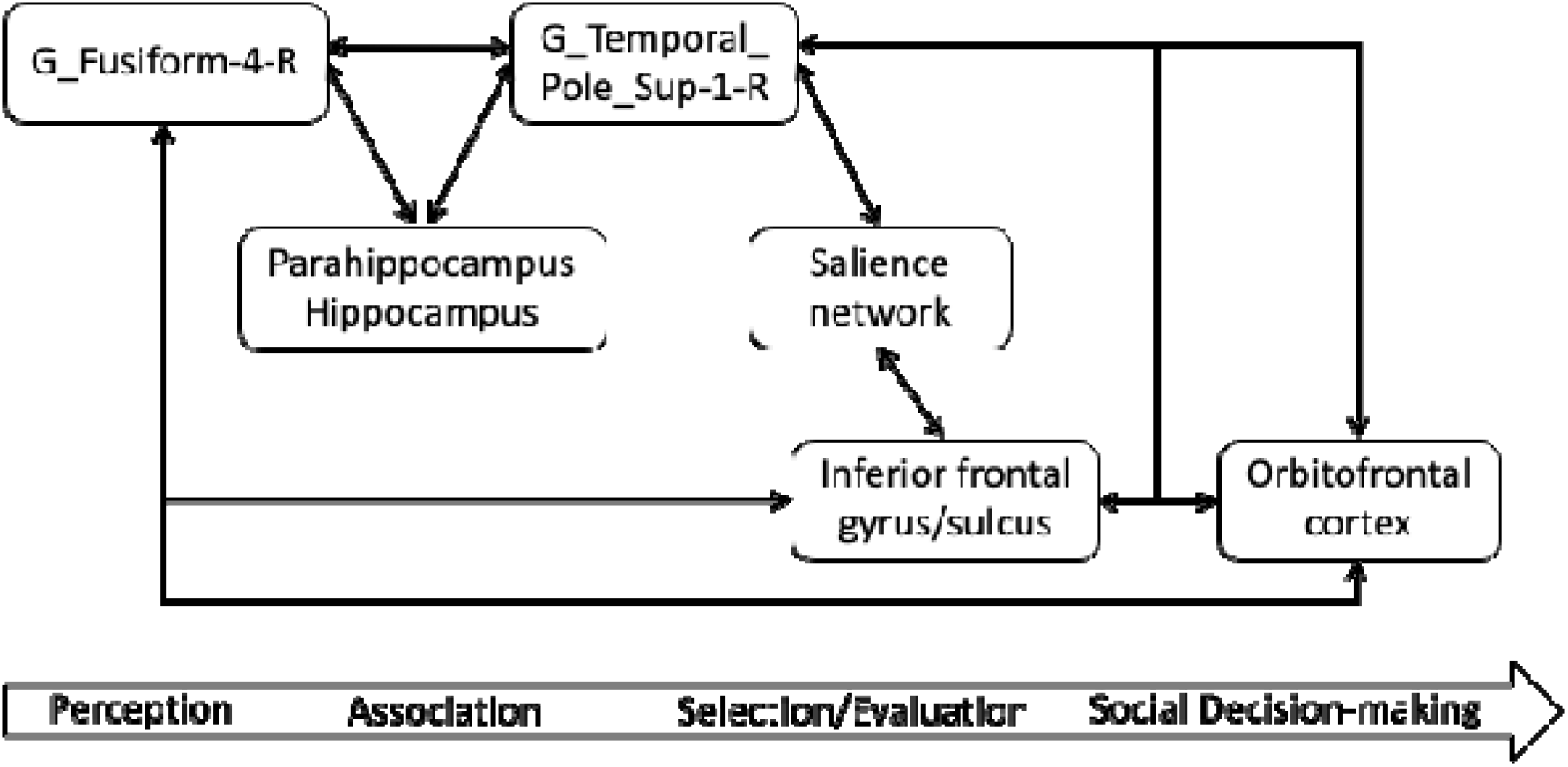

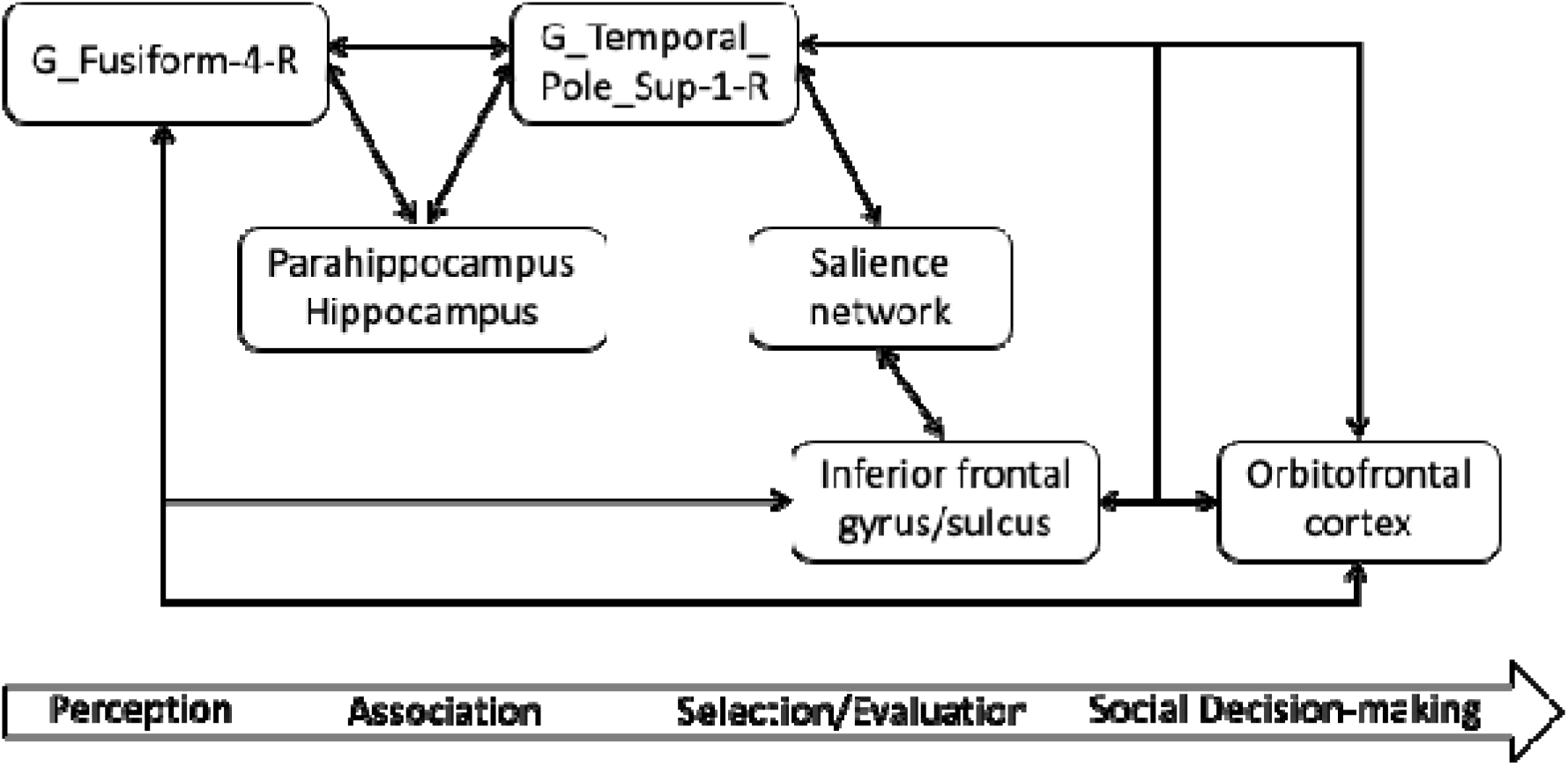

Notably, none of the regions involved in assigning social status exclusively dealt with social information. Most of these regions are involved in many cognitive functions. The functional connection of structures whose processing properties are beneficial for the execution of the task allows the social judgment function to emerge. Human connections exceed those of animals, including primates, at both the structural and functional levels ^113–115^. Thanks to creating functional connections (linking social cognition, memory, and executive functions), humans could use symbolic items and markings to signify social status.

## 5. Conclusion

This study delved into the neural mechanisms involved in the social interpretation of facial adornments and found that various brain regions, including the FFA, temporal poles, salience network, and orbitofrontal cortex, were involved in this process. Furthermore, assigning a social status from symbolic cues also activated the medial temporal regions and the inferior frontal gyrus, reflecting the role of episodic memory, contextual association, and executive functions. The complexity of this neural network raises questions about when it became fully functional in our ancestors and whether it resulted from a gradual process of integration and complexification or was already fully functional when the first archaeological evidence of culturalization of the human face was recorded.

The gradual complexification and patchy emergence of face adornment technologies over the last 500,000 years suggest a scenario of increasing but asynchronous integration of brain areas involved in social status recognition based on facial culturalization. This growing integration allowed the decoding of increasingly complex symbolic codes, supported by more demanding technologies for face adornment.

The interplay between cultural and biological mechanisms likely drove this process, with individuals gifted in acquiring, decoding, and creating these symbolic messages having selective advantages that favored the permanent inscription of a more integrated connectivity in the brain ^116–118^. A progressive co-option of brain regions has also been suggested for the evolution of tool-making ^119–121^. It would have enabled the development of increasingly complex tools.

The period between 140,000 and 70,000 years ago may have represented a key moment in this integration process, as this was when red pigments use became almost ubiquitous at African Middle Stone Age sites and marine shell beads were used for the first time in North Africa, the Near East, and Southern Africa. This diversification of colors, shapes, and technologies indicates a complexification of practices allowing wearers to use their faces to communicate information about their social role using more complex shared symbolic codes. It is reasonable to think that the human brain had largely equipped itself with the necessary connections to process and interpret these stimuli 70,000 years ago.

## 6. Acknowledgments

We thank the Ginesis Lab (GIN, Fealinks, Labcom Programme 2016, ANR 16LCV2-0006-01) for their help with data management and processing. We are also indebted to Violaine Verrecchia and Marc Joliot for their help in data analysis. Our thanks also go to Marie Guerlain, Annie Bardon-Lay, and her team members for their help with the face painting. Warm thanks also go to all those who agreed to participate in our experiments.

## 7. Authors’ contributions

MS, FE, SR, EM designed the study

MS, EM acquired the data

MS, EM analyzed the data

MS, FE, SR, EM wrote the article

## 9. Data Availability

Raw BOLD values for all the hROIs and contrasts are available as supplementary material

