## Supplemental Table 1 for "Assigning a social status from face adornments: an fMRI study"

| **hROI Social status - Ornament check  (* Specific to Social status – Ornament check)** | **MNI coordinates of the center of mass** | **Mean (BOLD)** | **Standard deviation (BOLD)** | **p (FDR corrected)** |
| --- | --- | --- | --- | --- |
| **G_Cingulum_Ant-2-L*** | -7 34 22 | 0.12 | 0.16 | 0.0005 |
| G_Cingulum_Ant-2-R | 7 33 23 | 0.08 | 0.16 | 0.0173 |
| G_Cingulum_Mid-2-L | -4 3 30 | 0.15 | 0.16 | <.0001 |
| G_Cingulum_Mid-2-R | 4 4 30 | 0.11 | 0.18 | 0.0045 |
| **S_Cingulate-1-L*** | -7 27 30 | 0.23 | 0.20 | <.0001 |
| S_Cingulate-1-R | 7 27 31 | 0.20 | 0.19 | <.0001 |
| S_Cingulate-2-L | -7 16 41 | 0.22 | 0.22 | <.0001 |
| S_Cingulate-2-R | 8 14 46 | 0.14 | 0.20 | 0.0011 |
| G_Cingulum_Post-1-L | -4 -26 29 | 0.12 | 0.22 | 0.0072 |
| G_Cingulum_Post-2-L | -4 -39 27 | 0.11 | 0.23 | 0.0121 |
| G_Cingulum_Post-3-L | -5 -43 10 | 0.13 | 0.24 | 0.0071 |
| G_Cingulum_Post-3-R | 6 -42 10 | 0.10 | 0.22 | 0.0253 |
| **G_Frontal_Inf_Orb-1-L*** | -42 31 -17 | 0,21 | 0.22 | <.0001 |
| **G_Frontal_Inf_Orb-1-R*** | 44 33 -14 | 0.13 | 0.16 | 0.0001 |
| **G_Frontal_Mid_Orb-2-L*** | -41 49 -5 | 0.23 | 0.30 | 0.0005 |
| S_Orbital-1-R | 25 41 -15 | 0.07 | 0.11 | 0.0023 |
| **S_Orbital-2-L*** | -31 34 -13 | 0.26 | 0.21 | <.0001 |
| **S_Orbital-2-R*** | 29 34 -13 | 0.18 | 0.17 | <.0001 |
| G_Frontal_Mid-1-R | 41 44 13 | 0.09 | 0.16 | 0.0075 |
| G_Frontal_Mid-5-L | -43 20 37 | 0.21 | 0.35 | 0.0038 |
| G_Frontal_Mid-5-R | 42 17 41 | 0.09 | 0.17 | 0.0132 |
| **S_Inf_Frontal-1-L*** | -44 38 12 | 0.37 | 0.31 | <.0001 |
| **S_Inf_Frontal-1-R*** | 46 40 10 | 0.24 | 0.21 | <.0001 |
| S_Inf_Frontal-2-L | -43 15 29 | 0.38 | 0.34 | <.0001 |
| **S_Inf_Frontal-2-R*** | 44 19 28 | 0.28 | 0.21 | <.0001 |
| **G_Frontal_Inf_Tri-1-L*** | -49 26 5 | 0.15 | 0.22 | 0.0011 |
| **G_Frontal_Inf_Tri-1-R*** | 50 29 5 | 0.07 | 0.17 | 0.0434 |
| S_Precentral-1-R | 50 10 24 | 0.11 | 0.24 | 0.0178 |
| **G_Frontal_Sup_Medial-3-L*** | -5 35 43 | 0.19 | 0.23 | 0.0002 |
| G_Frontal_Sup_Medial-3-R | 6 33 44 | 0.14 | 0.19 | 0.0006 |
| **G_Supp_Motor_Area-1-L*** | -6 22 46 | 0.39 | 0.23 | <.0001 |
| **G_Supp_Motor_Area-1-R*** | 6 21 48 | 0.37 | 0.26 | <.0001 |
| G_Supp_Motor_Area-2-L | -11 18 61 | 0.08 | 0.16 | 0.0134 |
| **G_Insula-anterior-1-L*** | -20 5 -19 | 0.12 | 0.18 | 0.0013 |
| **G_Insula-anterior-1-R*** | 19 7 -19 | 0.07 | 0.17 | 0.0329 |
| **G_Insula-anterior-2-L*** | -34 17 -13 | 0.17 | 0.24 | 0.0009 |
| G_Insula-anterior-2-R | 35 18 -13 | 0.10 | 0.18 | 0.0054 |
| **G_Insula-anterior-3-L*** | -34 24 1 | 0.31 | 0.22 | <.0001 |
| G_Insula-anterior-3-R | 37 24 0 | 0.20 | 0.15 | <.0001 |
| G_Insula-anterior-4-L | -41 15 3 | 0.08 | 0.21 | 0.0457 |
| G_Occipital_Inf-1-R | 50 -60 -9 | 0.22 | 0.19 | <.0001 |
| G_Occipital_Inf-2-L | -45 -71 -7 | 0.13 | 0.17 | 0.0003 |
| G_Occipital_Inf-2-R | 47 -65 -7 | 0.19 | 0.18 | <.0001 |
| G_Occipital_Lat-2-L | -26 -94 -1 | 0.06 | 0.15 | 0.0253 |
| G_Occipital_Lat-3-L | -40 -84 -12 | 0.10 | 0.16 | 0.0041 |
| G_Occipital_Lat-3-R | 43 -81 -10 | 0.11 | 0.20 | 0.0078 |
| G_Occipital_Lat-4-L | -31 -89 8 | 0.08 | 0.15 | 0.0078 |
| G_Occipital_Lat-4-R | 34 -85 9 | 0.09 | 0.15 | 0.0054 |
| G_Occipital_Lat-5-L | -35 -79 -1 | 0.07 | 0.15 | 0.0132 |
| G_Occipital_Lat-5-R | 36 -76 2 | 0.07 | 0.14 | 0.0116 |
| **G_Fusiform-2-L*** | -35 -26 -23 | 0.09 | 0.10 | <.0001 |
| G_Fusiform-4-L | -43 -50 -17 | 0.19 | 0.18 | <,0001 |
| **G_Fusiform-4-R*** | 44 -46 -18 | 0.22 | 0.15 | <.0001 |
| G_Fusiform-5-L | -31 -50 -12 | 0.09 | 0.12 | 0.0006 |
| G_Fusiform-5-R | 32 -47 -41 | 0.09 | 0.12 | 0.0006 |
| G_Fusiform-6-R | 29 -62 -9 | 0.05 | 0.12 | 0.0489 |
| S_Intraoccipital-1-L | -24 -72 32 | 0.09 | 0.23 | 0.0455 |
| S_Intraoccipital-1-R | 28 -69 33 | 0.15 | 0.21 | 0.0007 |
| **G_Precuneus-2-R*** | 5 -56 20 | 0.27 | 0.29 | <.0001 |
| **G_Precuneus-7-L*** | -6 -65 35 | 0.16 | 0.31 | 0.0097 |
| **G_Precuneus-7-R*** | 7 -63 36 | 0.21 | 0.29 | 0.0006 |
| S_Intraparietal-2-L | -34 -58 45 | 0.22 | 0.29 | 0.0005 |
| S_Intraparietal-2-R | 37 -52 48 | 0.14 | 0.20 | 0.0010 |
| S_Intraparietal-3-L | -27 -60 43 | 0.16 | 0.27 | 0.0051 |
| S_Intraparietal-3-R | 27 -61 46 | 0.13 | 0.19 | 0.0012 |
| G_Temporal_Inf-4-R | 54 -58 -11 | 0.10 | 0.19 | 0.0086 |
| **G_Temporal_Pole_Sup-1-L*** | -35 11 -24 | 0.09 | 0.13 | 0.0019 |
| **G_Temporal_Pole_Sup-1-R*** | 36 16 -24 | 0.10 | 0.16 | 0.0044 |
| **G_Temporal_Pole_Mid-2-L*** | -35 9 -33 | 0.05 | 0.10 | 0.0097 |
| **G_Temporal_Pole_Mid-2-R*** | 35 12 -34 | 0.04 | 0.09 | 0.0121 |
| **G_Hippocampus-1-L*** | -30 -7 -19 | 0.06 | 0.11 | 0.0115 |
| **G_Hippocampus-1-R*** | 30 -5 -18 | 0.08 | 0.11 | 0.0005 |
| G_Hippocampus-2-L | -25 -32 -3 | 0.04 | 0.09 | 0.0430 |
| G_Hippocampus-2-R | 25 -31 -2 | 0.06 | 0.09 | 0.0041 |
| **G_ParaHippocampal-1-L*** | -16 -4 -18 | 0.24 | 0.23 | <.0001 |
| **G_ParaHippocampal-1-R*** | 14 -4 -18 | 0.17 | 0.20 | 0.0001 |
| **G_ParaHippocampal-2-L*** | -28 -27 -19 | 0.07 | 0.17 | 0.0253 |
| G_ParaHippocampal-2-R | 29 -25 -19 | 0.07 | 0.14 | 0.0169 |
| **G_ParaHippocampal-4-L*** | -17 -27 -13 | 0.12 | 0.21 | 0.0071 |
| **G_ParaHippocampal-4-R*** | 17 -27 -10 | 0.15 | 0.18 | 0.0002 |
| N_Caudate-4-R | 14 20 8 | 0.06 | 0.13 | 0.0237 |
| N_Caudate-5-L | -13 10 8 | 0.17 | 0.21 | 0.0003 |
| N_Caudate-5-R | 12 10 9 | 0.16 | 0.22 | 0.0006 |
| N_Thalamus-1-L | -4 0 1 | 0.18 | 0.29 | 0.0032 |
| N_Thalamus-1-R | 4 0 1 | 0.16 | 0.24 | 0.0012 |
| N_Thalamus-2-R | 9 -7 13 | 0.09 | 0.18 | 0.0120 |
| N_Thalamus-3-L | -3 -7 -1 | 0.15 | 0.22 | 0.0017 |
| **N_Thalamus-4-L*** | -3 -14 8 | 0.20 | 0.27 | 0.0005 |
| N_Thalamus-4-R | 3 -14 9 | 0.17 | 0.25 | 0.0017 |
| N_Thalamus-5-L | -12 -19 7 | 0.09 | 0.14 | 0.0026 |
| N_Thalamus-5-R | 13 -17 6 | 0.07 | 0.14 | 0.0148 |
| N_Thalamus-6-R | 15 -27 13 | 0.05 | 0.12 | 0.0455 |
| N_Thalamus-7-L | -9 -28 11 | 0.09 | 0.15 | 0.0054 |
| N_Thalamus-9-L | -5 -11 -7 | 0.12 | 0.13 | <.0001 |
| N_Thalamus-9-R | 5 -10 -6 | 0.11 | 0.16 | 0.0007 |
