## Supplemental Table 2 for "Assigning a social status from face adornments: an fMRI study"

Table 1: p-value (FDR corrected) < 0.05 of the t-test on each connection between G_Fusiform-4-R and the 37 hROIs positive in contrast [Social status minus Ornament check] and positive in contrast [Ornament (1-back) minus Face].

| **G_Fusiform-4-R** | **hROI** | **p (FDR corrected)** |
| --- | --- | --- |
|  | G_Fusiform-2-L | <.0001 |
|  | G_Frontal_Inf_Orb-1-L | 0.0248 |
|  | G_Frontal_Inf_Tri-1-L | 0.0004 |
|  | G_Frontal_Inf_Tri-1-R | 0.0033 |
|  | S_Inf_Frontal-1-L | <.0001 |
|  | S_Inf_Frontal-1-R | <.0001 |
|  | S_Inf_Frontal-2-R | <.0001 |
|  | S_Orbital-2-L | 0.0006 |
|  | S_Orbital-2-R | <.0001 |
|  | G_Hippocampus-1-L | <.0001 |
|  | G_Hippocampus-1-R | <.0001 |
|  | G_Insula-anterior-1-L | 0.0019 |
|  | G_Insula-anterior-1-R | 0.0007 |
|  | G_ParaHippocampal-1-L | <.0001 |
|  | G_ParaHippocampal-1-R | 0.0001 |
|  | G_ParaHippocampal-2-L | <.0001 |
|  | G_ParaHippocampal-4-L | 0.0097 |
|  | G_ParaHippocampal-4-R | 0.0227 |
|  | G_Temporal_Pole_Mid-2-L | 0.0097 |
|  | G_Temporal_Pole_Mid-2-R | 0.0121 |
|  | G_Temporal_Pole_Sup-1-L | 0.0048 |
|  | G_Temporal_Pole_Sup-1-R | 0.0008 |

Table 2: p-value (FDR corrected) < 0.05 of the t-test on each connection between G_Temporal_Pole_Sup-1-R and the 37 hROIs positive in contrast [Social status minus Ornament check] and positive in contrast [Ornament (1-back) minus Face].

| **G_Temporal_Pole_Sup-1-R** | **hROI** | **p (FDR corrected)** |
| --- | --- | --- |
|  | G_Hippocampus-1-L | <.0001 |
|  | G_Hippocampus-1-R | <.0001 |
|  | G_Insula-anterior-1-L | <.0001 |
|  | G_Insula-anterior-1-R | <.0001 |
|  | G_Temporal_Pole_Sup-1-L | <.0001 |
|  | G_Temporal_Pole_Mid-2-R | <.0001 |
|  | G_Frontal_Inf_Orb-1-R | <.0001 |
|  | G_Insula-anterior-2-L | <.0001 |
|  | G_Frontal_Inf_Tri-1-R | <.0001 |
|  | G_Frontal_Inf_Orb-1-L | <.0001 |
|  | G_Temporal_Pole_Mid-2-L | <.0001 |
|  | G_Frontal_Inf_Tri-1-L | <.0001 |
|  | G_ParaHippocampal-2-L | <.0001 |
|  | G_ParaHippocampal-1-L | <.0001 |
|  | G_Cingulum_Ant-2-L | <.0001 |
|  | G_Precuneus-7-R | 0.0001 |
|  | G_ParaHippocampal-1-R | 0.0002 |
|  | G_Frontal_Sup_Medial-3-L | 0.0002 |
|  | G_Fusiform-4-R | 0.0008 |
|  | G_Precuneus-7-L | 0.0055 |
|  | G_Insula-anterior-3-L | 0.0060 |
|  | S_Orbital-2-R | 0.0064 |
|  | G_Fusiform-2-L | 0.0067 |
|  | G_ParaHippocampal-4-L | 0.0068 |
|  | G_Supp_Motor_Area-2-L | 0.0160 |
|  | G_Precuneus-2-R | 0.0221 |
|  | S_Inf_Frontal-2-R | 0.0421 |
